# Pp6-Pfkfb1 axis modulates intracellular bacterial proliferation by orchestrating host-pathogen metabolic crosstalk

**DOI:** 10.1101/2025.06.20.660667

**Authors:** Li Fan, Yang Sun, Fangzhou Lou, Zilong Fang, Wengxiang Ding, Xiangxiao Li, Yan Li, Qingqing Shen, Siyu Deng, Jihuan Liang, Fengjiao Zhang, Sibei Tang, Zhikai Wang, Xiaojie Cai, Jiajia Tong, Zhenyao Xu, Jing Zou, Qing Yang, Honglin Wang

## Abstract

Intracellular bacterial pathogens exhibit heterogeneous replication rates within host macrophages, but the mechanisms by which they manipulate host factors for survival remain incompletely understood. Using a fluorescence-dilution reporter system in *Salmonella typhimurium* (*Salmonella*)-infected macrophages, we found that Protein Phosphatase 6 (Pp6) was downregulated in macrophages harboring growing bacteria. Conditional knockout of *Pp6* elevated host susceptibility to *Salmonella*-mediated lethality due to compromised antimicrobial defenses. MicroRNA-31 (miR-31) was identified as a negative regulator of Pp6, and its conditional ablation enhanced bacterial clearance. Yeast two-hybrid screening identified 6-phosphofructo-2-kinase/fructose-2,6-biphosphatase 1 (Pfkfb1), a metabolic regulator, as a substrate of Pp6. *Pp6* deficiency resulted in significantly elevated expression of Pfkfb1, which was highly expressed in macrophages containing replicating *Salmonella*. *Pfkfb1* deletion restricted bacterial proliferation by promoting nitric oxide (NO) production while concurrently suppressing arginase-1 (Arg-1) expression and impairing arginine metabolism in macrophages. Collectively, these results establish the Pp6-Pfkfb1 axis as a key regulator of host metabolic adaptation and intracellular bacterial survival, offering potential therapeutic targets against multidrug-resistant pathogens.

**Author summary:** Intracellular pathogens secrete effector proteins that intercept and modify host cells to usurp host defenses and establish habitable intracellular niches, yet the host factors they manipulate for survival and virulence are not fully understood. Using a fluorescence-dilution reporter system in *Salmonella typhimurium* (*Salmonella*)-infected macrophages, we found Protein Phosphatase 6 (Pp6) was significantly decreased in macrophages harboring replicating *Salmonella* and its deficiency indulged bacterial growth in macrophages. However, the deletion of *microRNA-31* (*miR-31*), a negative regulator of Pp6, enhanced *Salmonella* clearance. Furthermore, through yeast two-hybrid screening, we identified 6-phosphofructo-2-kinase/fructose-2,6-biphosphatase 1 (Pfkfb1) is a substrate of Pp6. And we demonstrated Pfkfb1, abundantly expressed in macrophages containing growing bacteria, is crucial for *Salmonella* proliferation by regulating nitric oxide (NO) levels, arginase-1 (Arg-1) expression and arginine metabolism. Overall, the molecular changes in Pp6-Pfkfb1 axis drive host metabolic adaptations that enable intracellular bacterial survival.

## Introduction

Macrophages serve essential functions in preserving tissue homeostasis and providing host defense against pathogens [1]. Upon exposure to microbial signals, damaged tissue components, or activated lymphocyte-derived stimuli, macrophages undergo metabolic and functional reprogramming, resulting in distinct phenotypic states that dictate their capacity to regulate immune response [2]. When activated by lipopolysaccharide (LPS) or interferon (IFN)-γ, macrophages are polarized to a pro-inflammatory M1 phenotype, which harbor elevated intracellular nitric oxide (NO) levels and strong bactericidal activity. Under alternative conditions, interleukin-4 (IL-4) and interleukin-13 (IL-13) stimulation induces macrophage polarization toward a M2 phenotype, which exhibits increased arginase-1 (Arg-1) expression accompanied by metabolic reprogramming of arginine metabolism toward ornithine and polyamine biosynthesis. M2-polarized macrophages primarily mediate tissue remodeling and anti-inflammatory response [3–5].

*Salmonella typhimurium* (*Salmonella*) is an invasive, facultative enteric pathogen responsible for gastroenteritis and life-threatening systemic diseases in humans and animals [6]. Macrophages engulf intracellular bacterial pathogens, such as *Salmonella*, which subsequently replicates within host cells and causes difficult-to-eradicate infections. Once internalized, *Salmonella* triggers multiple host cell responses, including specific metabolic alterations mediated by certain virulence factors to provide energy sources and suitable environments for bacterial growth [7, 8]. Multiple bacteria-centered mechanisms required for intracellular survival have been unveiled. For instance, the ZnuABC transporter and its associated glycolytic activity in *Salmonella* contribute to antinitrosative defenses against NO [9]; reactive persulfides from *Salmonella* inhibit autophagy-mediated bacterial clearance in macrophages [10]; *Salmonella* Pathogenicity Island 2-encoded type III secretion system confers resistance to reactive oxygen species [6]. Nevertheless, host cellular components manipulated by *Salmonella* to indulge bacterial replication remain poorly defined. A recent single-cell transcriptomic analysis revealed that macrophages harboring non-growing *Salmonella* display hallmarks of pro-inflammatory M1 polarization, whereas those containing growing bacteria undergo metabolic reprogramming toward anti-inflammatory M2-like state [11, 12]. Therefore, deciphering the mechanistic basis underlying macrophage state transitions between non-growing or growing bacteria will uncover potential therapeutic targets and advance the treatment strategies for intracellular bacterial infections.

Here, our study demonstrates that during *Salmonella* infection, Protein Phosphatase 6 (Pp6) exhibits marked reduction within macrophages containing replicating bacteria and its absence facilitates *Salmonella* proliferation and shortens the survival of infected mice. In contrast, knockout of *microRNA-31* (*miR-31*), a negative Pp6 regulator, suppresses *Salmonella* replication. Critically, phosphofructo-2-kinase/fructose-2,6- biphosphatase 1 (Pfkfb1), an enzyme crucial for metabolism-oriented macrophage polarization [13], is identified as a substrate of Pp6 and distinctly expressed in macrophages containing growing pathogens. *Pp6* deficiency impairs Pfkfb1 dephosphorylation and increases Pfkfb1 expression levels. Genetic ablation of *Pfkfb1* increases NO production and downregulates Arg-1 expression as well as arginine biosynthesis and metabolism, thereby restricting *Salmonella* replication in infected cells. Collectively, these findings highlight the Pp6-Pfkfb1 axis as a pivotal regulator of host-pathogen interactions, indicating that therapeutic targeting of this pathway may potentiate host antibacterial defenses.

## Results

### The deficiency of *Pp6* indulges bacterial replication in macrophages

Given the significant heterogeneity in growth rates exhibited by intracellular bacterial pathogens within host macrophages [11], a *Salmonella* strain harboring a dual-reporter plasmid (pFCcGi) was developed to monitor bacterial replication following internalization [14]. This system utilizes constitutively expressed mCherry and conditionally expressed GFP, permitting differentiation of three distinct states: (i) actively dividing bacteria (increasing mCherry/stable GFP), (ii) non-replicating bacteria (stable dual fluorescence), and (iii) bystander macrophages (double-negative) [11, 12] (**Fig 1A**). Fluorescence-activated cell sorting (FACS) successfully resolved the three macrophage populations: bystander, and those containing growing/non-growing bacteria (**Fig 1B**). We further validated the gating strategy effectively isolated macrophages with distinct intracellular bacterial loads by reanalyzing the sorted cells using fluorescence microscopy (**Fig 1B**). Immunocytochemistry analysis revealed a diminished level of Pp6 in macrophages harboring growing *Salmonella*, relative to non-infected controls, bystander macrophages and macrophages containing non-replicating bacteria (**Fig 1C**). This observation received further validation via western blot analysis (**Fig 1D**).

**Fig 1.**
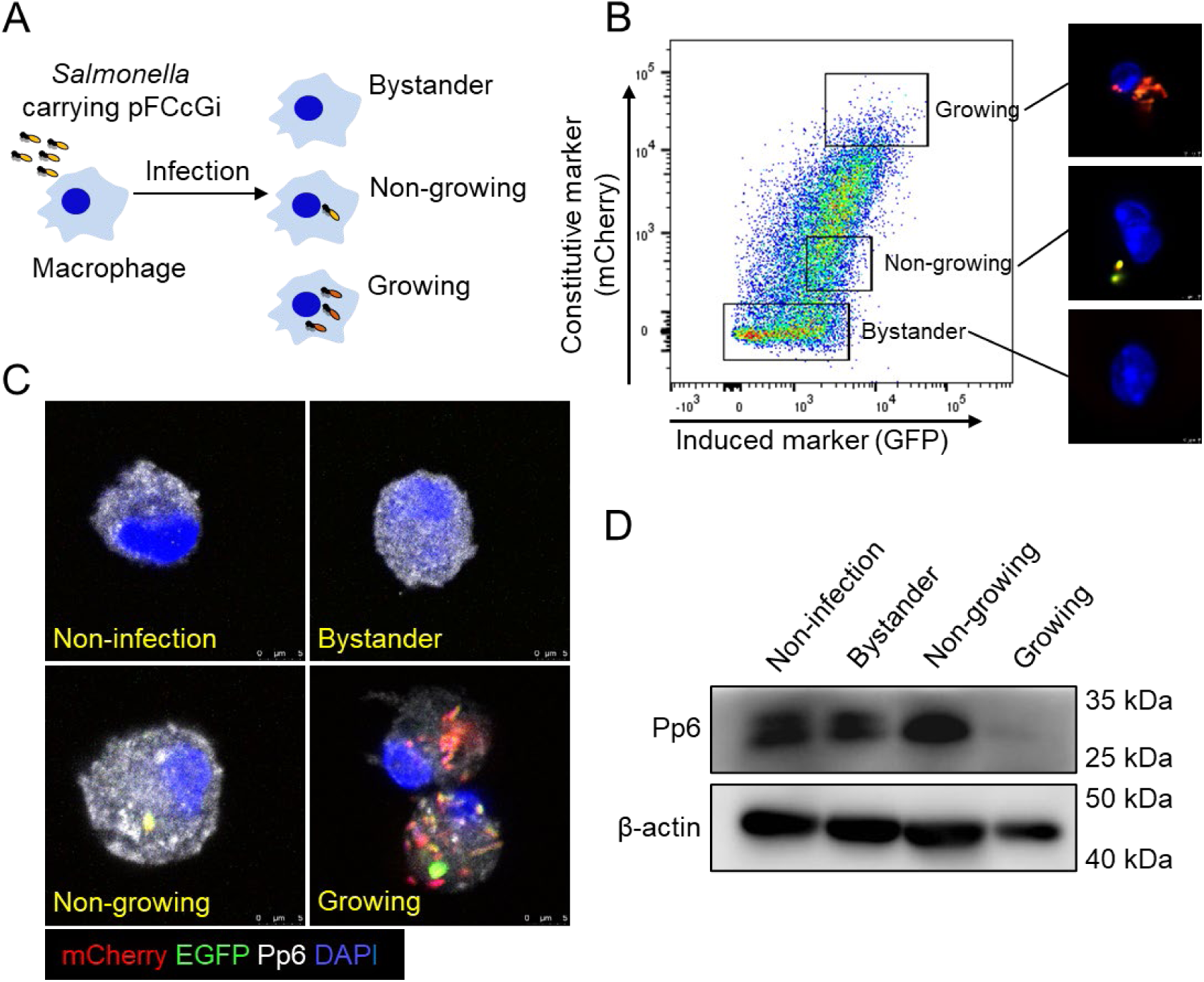
Pp6 expression is downregulated in macrophages harboring replicating *Salmonella*. (A) Experimental workflow of BMDMs infected with *Salmonella* carrying the dual-reporter plasmid (pFCcGi). Three distinct macrophage populations were identified: (i) uninfected bystanders, (ii) BMDMs with non-growing bacteria, and (iii) BMDMs with growing bacteria. (B) The populations were validated using FACS and further characterized through immunocytochemistry imaging. Scale bar, 3 μm. **(C-D)** Different cell subpopulations were isolated at 18 h post infection (hpi) and the expression of Pp6 was respectively detected in non-infected, bystander, non-growing and growing group by immunocytochemistry (C) and western blot (D). Scale bar, 5 μm.

As a member of the type 2A serine/threonine (Ser/Thr) protein phosphatase family, Pp6 is known to mediate diverse cellular functions through dynamic interactions with regulatory subunits and substrates, modulating processes such as cell cycle progression, DNA damage response, and immune signaling [15]. Despite this broad regulatory potential, the functional contributions of Pp6 to macrophage-mediated antimicrobial responses, particularly concerning intracellular bacterial pathogen, remain insufficiently characterized. To investigate the role of Pp6 in bacterial replication, *LysM*^Cre^*Pp6*^fl/fl^ mice were generated (**S1A Fig**), with macrophage-specific Pp6 depletion confirmed by western blot (**S1B Fig**). *In vitro* infection assays using bone marrow-derived macrophages (BMDMs) demonstrated significantly elevated bacterial loads in *LysM*^Cre^*Pp6*^fl/fl^ macrophages compared to *Pp6*^fl/fl^ controls during the replicative phase of pFCcGi-expressing *Salmonella*, as quantified by flow cytometry **(Figs 2A and B)**. For *in vivo* assessment, intraperitoneal challenge with a lethal dose of wild-type (WT) *Salmonella* resulted in markedly decreased survival in *LysM*^Cre^*Pp6*^fl/fl^ mice compared to *Pp6*^fl/fl^ mice **(Fig 2C)**. Whole-mount imaging of liver tissues at 3 days post-infection enabled spatial mapping of bacterial dissemination **(Fig 2D)**, revealing substantially enhanced bacterial colonization within the livers of *LysM*^Cre^*Pp6*^fl/fl^ mice versus littermate controls **(Figs 2E–H)**. These results collectively establish Pp6 as a critical host regulatory factor limiting intracellular bacterial proliferation.

**Fig 2.**
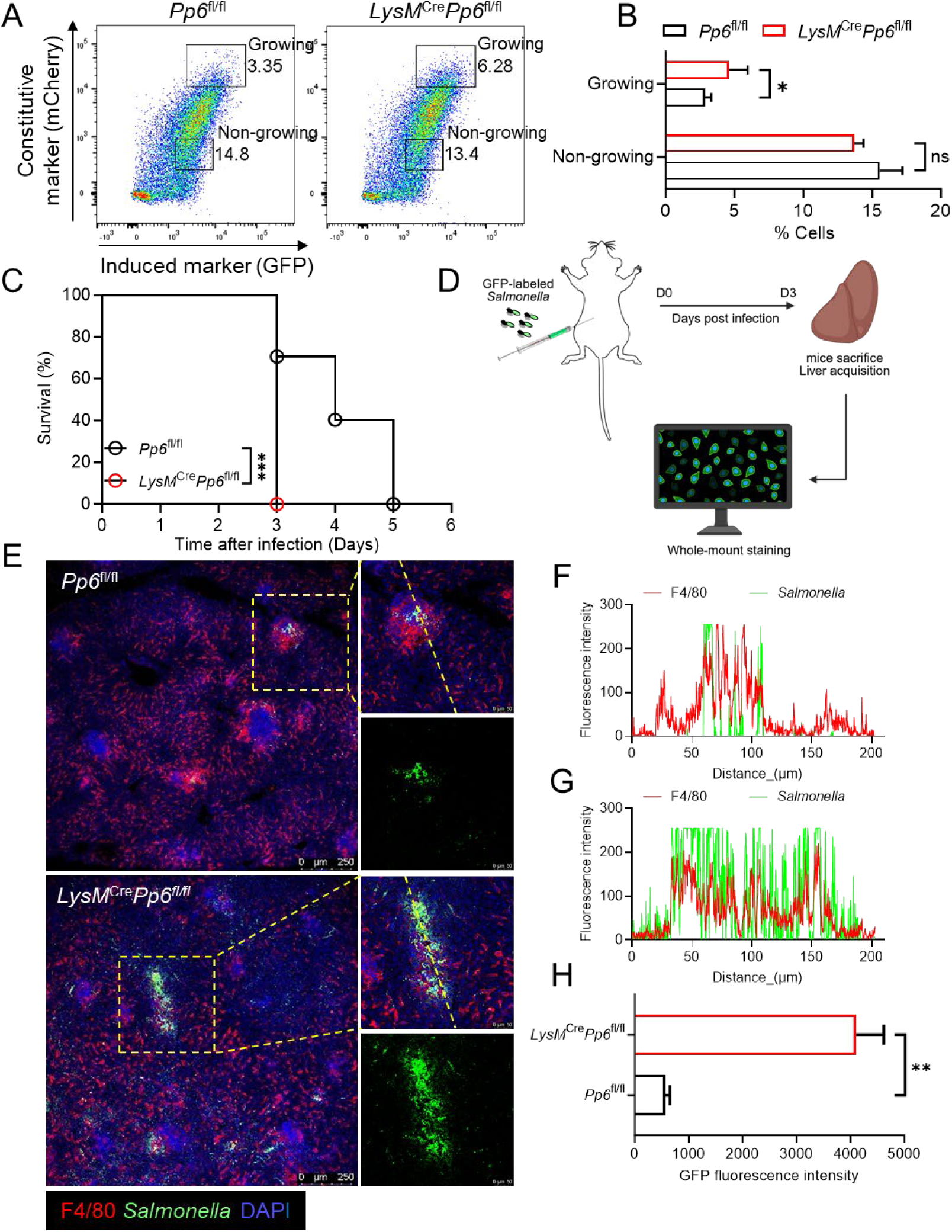
*Pp6* deficiency promotes intracellular hyperproliferation of *Salmonella* in macrophages. **(A-B)** Flow cytometry analysis of *Salmonella* proliferation in BMDMs from *LysM*^Cre^*Pp6*^fl/fl^ mice and *Pp6*^fl/fl^ mice (n = 4) (A); statistical data were shown in (B). *P*-values were determined by two-tailed unpaired *t*-test (mean ± SEM). ns, not significant, \**P* < 0.05. (C) Kaplan-Meier survival curves of *LysM*^Cre^*Pp6*^fl/fl^ mice and *Pp6*^fl/fl^ mice (n = 10) infected intraperitoneally with 1 = 10^3^ colony-forming units (CFU) WT *Salmonella*. *P*- value was determined by Log-rank (Mantel-Cox) test. \*\*\**P* < 0.001. (D) Schematic illustration of whole-mount staining of the mouse liver following *in vivo* infection. **(E-H)** Representative fluorescence labeling of GFP-labeled *Salmonella* and F4/80 in liver tissues from *LysM*^Cre^*Pp6*^fl/fl^ mice and *Pp6*^fl/fl^ mice following 3-day infection (E). The colocalization fluorescence intensity on the yellow dashed line of zoom in images (F, G) and the GFP signal intensity (H) were quantified using ImageJ software. Scale bar, 250 μm or 50 μm. *P*-value was determined by two-tailed unpaired *t*-test (mean ± SEM). \*\**P* < 0.01.

### *miR-31* deficiency rescues the expression of Pp6 and facilitates *Salmonella* clearance

Previous work identified that Pp6 as a key target of miR-31 in keratinocytes, with *miR- 31* deletion restoring the expression of Pp6 [16]. Therefore, we constructed a luciferase-expressing plasmid to investigate the post-transcriptional regulation mediated by *miR- 31* binding to the 3’ untranslated region (UTR) of *Pp6* mRNA in macrophages (**Fig 3A**). As a result, forced *miR-31* expression in RAW264.7 cells significantly reduced luciferase activity driven by the WT 3’ UTR plasmid, whereas no reduction occurred with the mutated plasmid lacking the target sequence (**Fig 3B**). This confirmed direct targeting of *Pp6* by *miR-31* in macrophages. Measurement of Pp6 protein levels in BMDMs from *miR-31*^-/-^ mice and littermate controls at different time points after infection revealed progressive Pp6 reduction from 2 to 12 hours in WT controls. Conversely, *miR-31*^-/-^ macrophages exhibited partial Pp6 restoration during this period (**Fig 3C**). We next generated *LysM*^Cre^*miR-31*^fl/fl^ mice by crossing *miR-31* floxed mice with *LysM* Cre transgenic mice [16, 17] (**S2A and B Figs**). Infection of BMDMs from *LysM*^Cre^*miR-31*^fl/fl^ and *miR-31*^fl/fl^ mice with *Salmonella* carrying pFCcGi revealed a decreased proportion of *LysM*^Cre^*miR-31*^fl/fl^ macrophages containing replicating bacteria in compared to *miR-31*^fl/fl^ controls (**Figs 3D and E**). Moreover, following intraperitoneal challenge with a lethal dose of WT *Salmonella*, *LysM*^Cre^*miR-31*^fl/fl^ mice exhibited a significantly enhanced survival rate than controls (**Fig 3F**). To further assess bacterial proliferation in macrophages, we infected mice with GFP-labeled *Salmonella* and observed a significantly reduced colonization in *LysM*^Cre^*miR-31*^fl/fl^ mice at day 3 after *Salmonella* infection (**Figs 3G-J**). Taken together, our findings demonstrate that *miR-31* deletion rescues Pp6 expression, reduces the proportion of macrophages harboring growing *Salmonella*, and enhances host resistance to infection.

**Fig 3.**
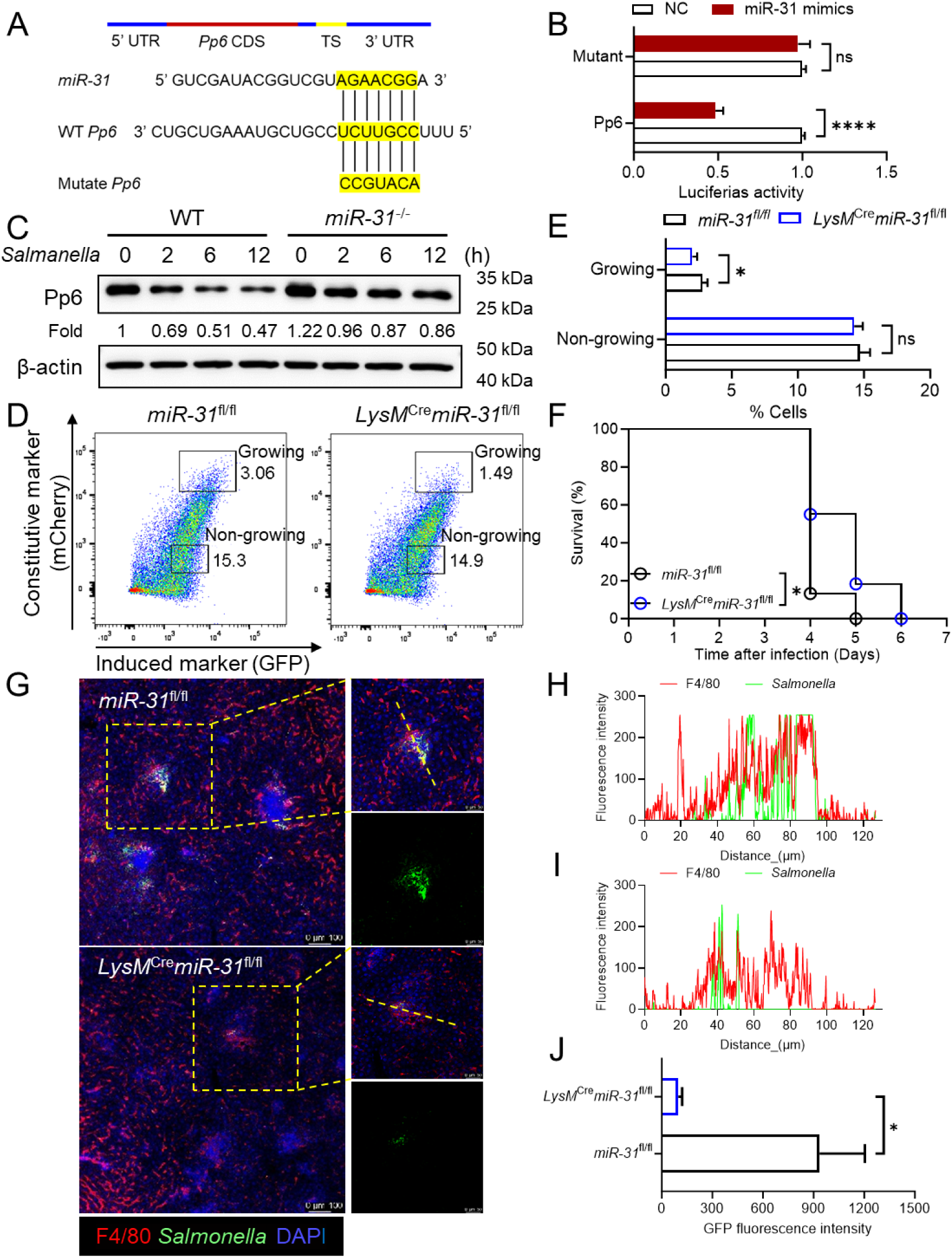
*miR-31* deficiency impairs *Salmonella* proliferation in macrophages. (A) WT and point-mutated 3’ UTR reporter constructs. (B) Luciferase activity assessment in RAW264.7 cells transfected with *miR-31* mimics and either the WT or point-mutated 3′ UTR reporter construct (designated as WT UTR or mutant UTR). *P*-values were determined by two-tailed unpaired *t*-test (mean ± SEM). ns, not significant, \*\*\*\**P* < 0.0001. (C) Western blot analysis of Pp6 protein levels in BMDMs from WT and *miR-31*^-/-^ mice. **(D-E)** Flow cytometry analysis of infected BMDMs from *LysM*^Cre^*miR-31*^fl/fl^ and *miR- 31*^fl/fl^ mice (n = 4) (D). And statistics of differential cell groups with non-growing or growing bacteria (E). *P*-values were determined by two-tailed unpaired *t*-test (mean ± SEM). ns, not significant, \**P* < 0.05. **(F)** Kaplan-Meier survival curves of *LysM*^Cre^*miR-31*^fl/fl^ and *miR-31*^fl/fl^ mice (n = 14) infected intraperitoneally with 1 = 10^3^ CFU WT *Salmonella*. *P*-value was determined by Log-rank (Mantel-Cox) test. \**P* < 0.05. **(G-J)** Representative whole-mount fluorescence images of GFP-labeled *Salmonella* and F4/80 in liver tissues from *LysM*^Cre^*miR-31*^fl/fl^ mice and *miR-31*^fl/fl^ mice 3 days post- infection (G). The colocalization fluorescence intensity on the yellow dashed line of zoom in images (H, I) and the GFP signal intensity (J) were quantified using ImageJ software. Scale bar, 100 μm or 50 μm. *P*-values were determined by two-tailed unpaired *t*-test (mean ± SEM). \**P* < 0.05.

### Pfkfb1 is a substrate of Pp6

To identify Pp6 substrates critical for macrophage functional regulation, a high-quality mouse universal cDNA library was employed in a yeast two-hybrid screening using Pp6 as bait. This approach identified 95 unique candidate substrates, with Pfkfb1 exhibiting robust binding affinity for Pp6 (**S1 Table and** **Fig 4A**). Co-immunoprecipitation assays performed in 293FT cells expressing myc-tagged PP6 further validated the PP6-PFKFB1 interaction (**Figs 4B and C**). CRISPR/Cas9- mediated *PP6* ablation in 293FT cells significantly increased PFKFB1 protein expression (**Fig 4D**). To investigate the PP6-regulated phosphorylation sites on PFKFB1, LC-MS/MS analysis of PFKFB1 immunoprecipitates from WT and *PP6*- deficient 293FT cells was performed. The results detected significantly enhanced phosphorylation at Ser-2, Ser-33, and Ser-178 residues in *PP6*-depleted cells **(S3A-C Figs)**. Subsequent evaluation of PFKFB1 protein levels following alanine (Ala) substitution mutations at these phosphorylation sites revealed that PFKFB1^S33-mut^ and PFKFB1^S178-mut^ maintained expression patterns consistent with the control group **(Fig 4E)**. In contrast, the PFKFB1^S2-mut^ mutation abolished *PP6*-mediated regulation of PFKFB1 protein expression **(Fig 4E).** These findings identify PFKFB1 as a primary substrate of PP6 and establish that PP6 governs PFKFB1 protein stability through site-specific dephosphorylation at Ser-2.

**Fig 4.**
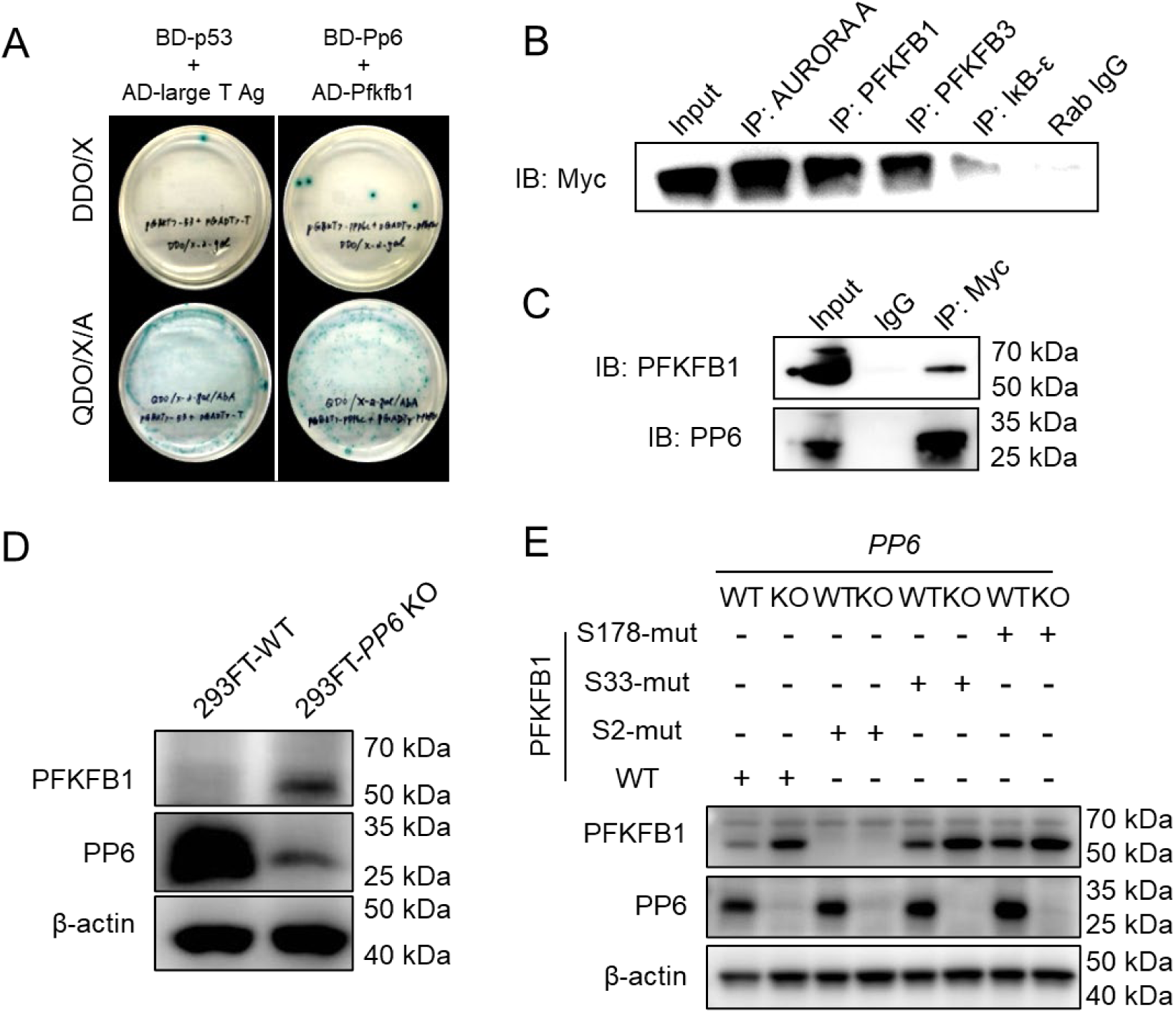
PFKFB1 is a substrate of PP6 (A) Identification of Pp6-interacting protein using the yeast two-hybrid system. (B) Cell lysates from 293FT cells expressing myc-tagged PP6 were immunoprecipitated with antibodies against AURORA A, PFKFB1, PFKFB3, IκB-ε, or rabbit IgG control, and then incubated with anti-Myc. AURORA A [29] and IκB-ε [30] have been reported to interact with PP6, serving as positive controls in this study. (C) Lysates were immunoprecipitated with IgG or anti-Myc, and blotted with anti- PFKFB1 and anti-PP6. (D) Endogenous *PP6* removal in 293FT cells using the CRISPR/Cas9 system and target protein expression analysis by western blot. (E) The phosphorylation sites of Ser-2, Ser-33 and Ser-178 were mutated with Ala substitution. Target protein levels were measured in WT and *PP6*-deficient 293FT cells transfected with these mutant plasmids.

### Pfkfb1 is essential for *Salmonella* growth and influences macrophage polarization

To determine whether Pfkfb1 functionally regulates intracellular bacterial replication within macrophages, Pfkfb1 expression was first assessed across FACS-sorted macrophage subsets: non-infected cells, infected cells harboring growing or non-growing bacteria, and bystander cells. Notably, Pfkfb1 expression was exclusively detected in macrophages containing replicating bacteria, concomitant with reduced Pp6 levels **(Fig 5A)**. Subsequently, we generated a *Pfkfb1* knockout (*Pfkfb1*^-/-^) mouse model to further explore its role **(S4A and B Figs)**. Infection of WT and *Pfkfb1*^-/-^ macrophages with pFCcGi-expressing *Salmonella* mediated a pronounced reduction in bacterial load within *Pfkfb1*^-/-^ macrophages relative to WT controls **(Figs 5B and C)**. Furthermore, intraperitoneally with a lethal dose of *Salmonella* resulted in significantly increased survival of *Pfkfb1*^-/-^ mice compared to WT littermate controls **(Fig 5D)**.

**Fig 5.**
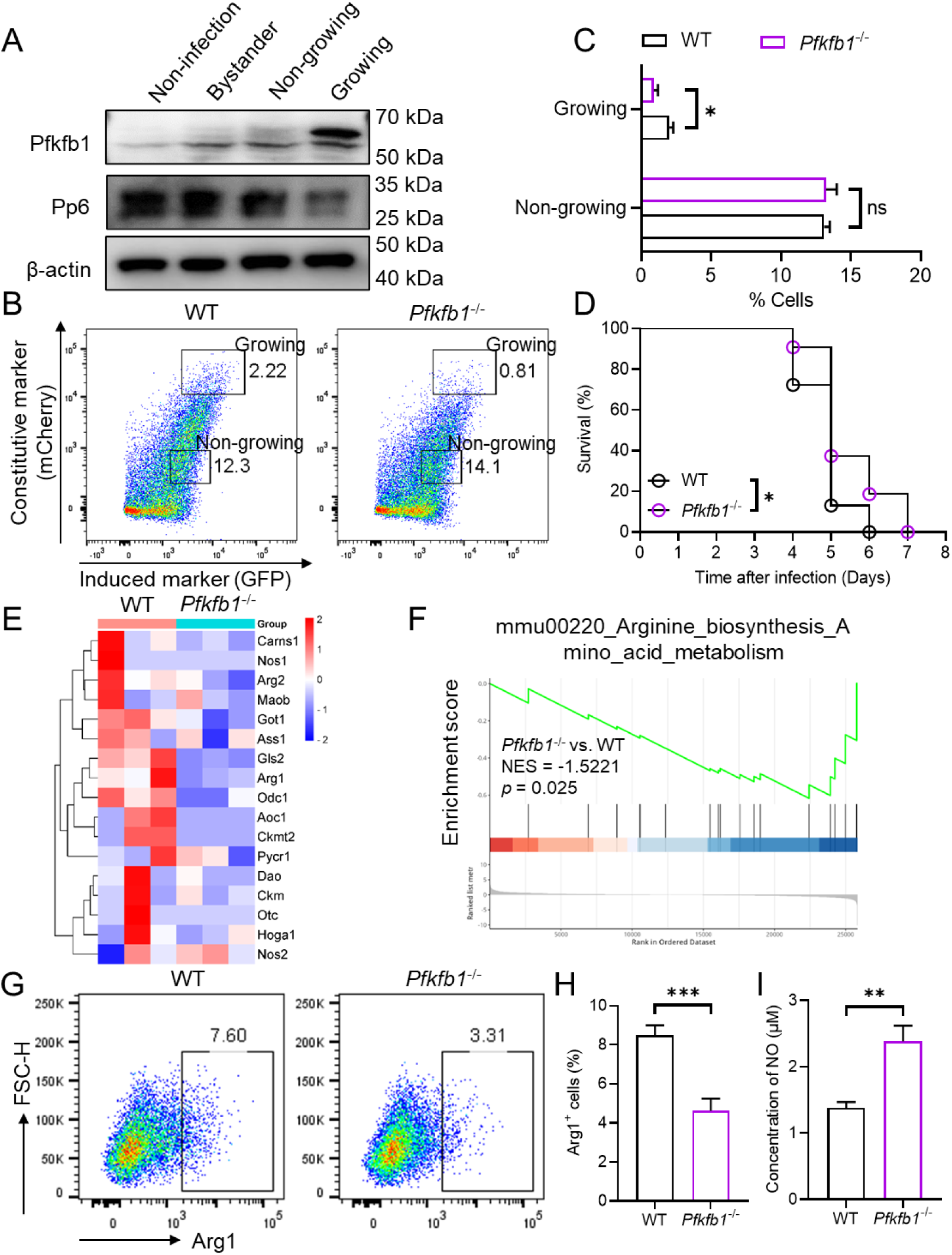
Pfkfb1 promotes bacteria growth and regulates macrophage polarization **(A)** Expression of Pfkfb1 and Pp6 in non-infected BMDMs, bystander BMDMs and BMDMs containing growing and non-growing bacteria was detected by western blot. **(B-C)** Flow cytometry analysis of *Salmonella* replication in BMDMs from WT and *Pfkfb1*^-/-^ mice (n = 4-5) (B). And statistics of differential cell groups with non-growing or growing bacteria (C). *P*-values were determined by two-tailed unpaired *t*-test (mean ± SEM). ns: not significant, \**P* < 0.05. (D) Kaplan-Meier survival curves of WT and *Pfkfb1*^-/-^ mice (n = 16-18) infected intraperitoneally with 1 = 10^3^ CFU WT *Salmonella*. *P*-value was determined by Log- rank (Mantel-Cox) test. \**P* < 0.05. (E) Heatmap of differentially expressed genes in arginine biosynthesis and metabolism (n = 3). (F) GSEA of differentially expressed genes in mouse BMDMs with arginine biosynthesis-associated genes enriched (n = 3). **(G-H)** Flow cytometry analysis (F) and quantification (G) of Arg-1 in WT and *Pfkfb1*^-/-^ BMDMs (n = 6). *P*-value was determined by two-tailed unpaired *t*-test (mean ± SEM). \*\*\**P* < 0.001. **(I)** The NO levels in WT and *Pfkfb1*^-/-^ BMDMs were measured using Griess regents (n = 6). *P*-value was determined by two-tailed unpaired *t*-test (mean ± SEM). \*\**P* < 0.01.

To explore the mechanistic basis of these findings, RNA-seq analysis was performed on WT *Salmonella*-infected BMDMs derived from WT and *Pfkfb1*^-/-^ mice. Interestingly, we found most genes implicated in arginine biosynthesis and metabolism exhibited reduced expression in BMDMs from *Pfkfb1*^-/-^ mice compared to WT controls **(Fig 5E)**. Gene set enrichment analysis (GSEA) further revealed the diminished activity in arginine biosynthesis pathway within *Pfkfb1*^-/-^ macrophages **(Fig 5F)**. Arginine serves as a common substrate for both arginase and nitric oxide synthase (NOS). Through flow cytometry analysis, we observed the expression of arginase-1 (Arg1), a characteristic marker of M2 macrophages, was markedly reduced in *Pfkfb1*^-/-^ macrophages **(Figs 5G and H)**. Concurrent assessment of NO expression levels demonstrated a marked increase in NO production in *Pfkfb1*^-/-^ cells relative to WT counterparts **(Fig 5I)**. Collectively, these results highlight Pfkfb1 as a critical metabolic regulator that modulates macrophage functionally and orchestrates host antimicrobial defense against bacterial infections.

## Discussion

Macrophage polarization is a dynamic process driven by extensive metabolic reprogramming, with the balance between pro-inflammatory M1 and anti-inflammatory M2 phenotypes playing a critical role in immune response [2, 4]. Although heterogeneity in macrophage polarization and *Salmonella* growth rate has been extensively investigated [11, 12, 18], the underlying mechanisms remain incompletely characterized. This study demonstrated that the host Pp6/Pfkfb1 axis drives macrophage polarization toward to M2 phenotype, creating a permissive niche for *Salmonella* survival and replication within macrophages. These findings suggest a potential therapeutic target for infectious diseases mediated by intracellular multidrug-resistant bacteria.

Pp6 exhibits high conservation across all eukaryotic species from yeast to humans, highlighting its fundamental biological significance. Mutations in Pp6 exist in 9-12.4% of melanomas and potentially drive melanoma development [19, 20]. Sit4/Ppe1, a Pp6 yeast homologue, is required for G1/S progression [21]. Human Pp6 plays critical roles in DNA damage responses via modulating DNA-dependent protein kinase (DNA-PK) signaling and homologous recombination-mediated repair of DNA double-strand breaks (DSBs) [22, 23], and interacts with Aurora A kinase [24]. Furthermore, recent studies indicate a broader role of Pp6 in pre-mRNA splicing [25], apoptosis regulation in immune cells [26], and Hippo pathway signaling modulation [27]. Collectively, these studies reveal that Pp6 integrates multiple signaling pathways. Current literature, however, indicates no prior investigation of the intrinsic function of Pp6 in bacterial replication within infected macrophages. Here, our research provides the first mechanistic evidence for Pp6-mediated regulation of intracellular bacterial growth in macrophages. We demonstrate that Pp6 expression is downregulated in macrophages during *Salmonella* infection, and that conditional deletion of *Pp6* enhances intracellular bacterial growth and significantly reduces host survival. These results establish Pp6 as a crucial regulator of immune responses against intracellular bacterial infections.

Previous study reported that miR-31 directly targets Pp6, with *miR-31* knockout rescuing the expression of Pp6 [16]. This regulatory relationship was confirmed within macrophages. Using genetic knockout mouse models, we further show that conditional ablation of *miR-31* attenuates intracellular bacterial proliferation in both *in vitro* and *in vivo* settings, thereby validating the functional role of Pp6 in macrophages during *Salmonella* infection.

Pfkfb1, expressed in M2 macrophages [28], remains poorly understood in the context of intracellular bacterial infections. Here, for the first time we identify Pfkfb1 is a direct substrate of Pp6, and *Pp6* deficiency increases Pfkfb1 expression and phosphorylation levels. Notably, Pfkfb1 expression is specifically detected in macrophages containing replicating bacteria. Experimental evidence confirms significant reduction in *Salmonella* growth and dissemination in *Pfkfb1*-deficient macrophages. Furthermore, Pfkfb1 is shown to regulate macrophage polarization through modulating NO and Arg- 1 expression, thereby controlling bacterial replication. While these findings establish involvement of Pfkfb1 in bacterial pathogenesis, the underlying mechanisms governing Pfkfb1-mediated regulation of intracellular bacterial growth require further studies.

In summary, this study reveals that Pp6 scavenges *Salmonella* from infected macrophages by negatively regulating Pfkfb1. During infection, Pfkfb1 regulates macrophages metabolic reprogramming to support bacteria proliferation within host cells. This regulatory mechanism represents a potential therapeutic target for interventions against infectious diseases caused by intracellular bacterial pathogens.

## Materials and Methods

### Bacterial strains and bacteria culture

The *Salmonella* strains used in this study were wild-type (WT) *Salmonella Typhimurium* S14028 (ATCC) strains along with two genetically modified derivatives: *Salmonella* carrying pFCcGi and GFP-labeled *Salmonella*. WT strains were cultured for 16 h at 37 °C in LB medium. The pFCcGi-carrying strains were cultured for 16 h at 37 °C in a MgMES pH 5.0 minimum medium containing 5 mM KCl, 7.5 mM (NH4)_2_SO_4_, 1 mM KH_2_PO_4_, 8 mM MgCl_2_, 38 mM glycerol, 0.1 % (w/v) casamino acids, 0.5 mM K_2_SO_4_, 170 mM MES pH 5.0 in distilled water (dH_2_O), supplemented with 0.2% L-arabinose and 100 μg/mL ampicillin. GFP-labeled *Salmonella* was cultured for 16 h at 37 °C in LB medium, supplemented with 15 μg/mL Gentamycin. Before infecting cells, bacteria were opsonized with 10% normal mouse serum (YEASEN) for 20 min at room temperature to obtain a phagocytosis rate of one bacterium per macrophage.

### Cell culture

Primary murine bone marrow-derived macrophages (BMDMs) were obtained from femoral and tibial bones of C57BL/6 WT mice or transgenic mice aged 8-10 weeks-old. The cells were cultured for 7 days in DMEM/high glucose medium (HyClone) containing 10% fetal bovine serum (FBS) (Gibco), 55 μM 2-Me (Gibco), 1% Penicillin/streptomycin/Amphotericin B (Solarbio), 100 μM Non-Essential amino acids (Solarbio), 10 mM HEPES (Solarbio), 1 mM sodium pyruvate (Solarbio) and 20 ng/mL macrophage colony-stimulating factor (M-CSF) (PeproTech), under standard culture conditions (37℃, 5% CO_2_). The culture medium was partially refreshed every 2 days. RAW264.7 cells were cultured in RPMI-1640 medium (Gibco) containing 5% FBS. HEK293FT cells were cultivated in DMEM/high glucose medium containing 10% FBS.

### Mice

C57BL/6 mice were purchased from Shanghai SLAC Laboratory Animal Co. Ltd. (Shanghai, China). *LysM* cre mice (stock number: 004781) were obtained from The Jackson Laboratory. The *miR-31*^fl/fl^ mice were generated as previously reported[16] and subsequently crossed with *LysM* cre transgenic mice. *Pp6*^fl/fl^ transgenic mice, provided by Dr. Wufan Tao, were similarly bred with *LysM* cre mice. The *Pfkfb1*^+/-^ mice were procured from Cyagen Biosciences. All experimental mice were maintained under specific pathogen-free (SPF) conditions in compliance with the National Institutes of Health Guide for the Care and Use of Laboratory Animals with the approval (SYXK- 2003-0026) by the Scientific Investigation Board of Shanghai Jiao Tong University School of Medicine (Shanghai, China).

### *In vitro* infection

To identify replicating and non-replicating intracellular *Salmonella*, BMDMs were plated at a density of 1.5x10^6^ cells per well in 12-well plate. Following overnight culture, the cells were infected with *Salmonella*-pFCcGi at a multiplicity of infection (MOI) of approximately 5 after washed with PBS. Centrifuge plate at 1600rpm for 5 min, followed by 30 min incubation at 37°C with 5% CO_2_. After incubation, the infected cells were washed three times with PBS and supplemented with fresh medium containing 100 μg/mL gentamicin for 30 min to eliminate extracellular bacteria. BMDMs were washed three times and incubated in medium with 5 μg/mL gentamicin for the remainder of the infection. After 18 h (Pp6 mice groups) or 22 h (other mice groups) of infection, cells were detached with 0.5% trypsin-EDTA (Gibco), stained with DAPI (BD Biosciences, 1:1000) for 10 min, and analyzed by flow cytometry (BD Fortessa) or sorted (BD FACS Aria III). FlowJo software was employed for data analysis.

### FACS sorting

To discriminate different subsets of BMDMs with bacteria carrying PFCcGi plasmid, cells were sorted post-infection. All experimental analyses excluded apoptotic macrophages and cellular doublets through appropriate gating strategies. Cell populations were isolated using a BD FACS Aria III sorter. Subsequently, sorted samples were processed for either western blot or immunofluorescence assay.

### Immunofluorescence

Infected-BMDMs were sorted into three groups according to intracellular bacterial fluorescence intensity. Each subpopulation was plated separately into 10 mm culture dishes. Bacterial contents were confirmed using fluorescence microscopy. Cells were washed with PBS and fixed with 4% paraformaldehyde for 20 min at room temperature (RT). Subsequent processing included two washes with 0.1% PBST, followed by incubation with primary anti-PPP6C antibody (Abcam, 1:100) for 2 h at RT. After two additional 0.1% PBST washes and one PBS wash, samples were incubated with fluorochrome-conjugated secondary antibodies (Life Technologies, 1:500) for 1 h at RT in dark. Then incubated with DAPI (BD Biosciences, 1:1000) for 5 min at RT in dark. Following two final 0.1% PBST washes and one PBS wash, samples were mounted using fluorescent mounting medium (Dako cat. S3023) and visualized with a Zeiss Leica TCS SP8 confocal microscope.

### *In vivo* infection

For survival rate assessment, WT *Salmonella* infection was established by infecting 8- 9 weeks-old mice with 1 × 10^3^ CFU bacteria administered intraperitoneally. Mortality was recorded daily. To monitor *in vivo* bacterial proliferation, GFP-labeled *Salmonella* infection was established by infecting 8-9 weeks-old mice with 1 × 10^3^ CFU bacteria administered intraperitoneally. At 3 days post-infection, animals were euthanized and liver tissues were collected for whole-mount immunofluorescence. For these experiments, mice were maintained in the P2 laboratory of Prof. X. Jia Group, Clinical Medicine Scientific and Technical Innovation Park, Shanghai Tenth Peoples Hospital

### Whole-mount immunofluorescence

Whole-mount samples were prepared following the Rapiclear® 1.52 Solution protocol. Briefly, C57BL/6 mice were euthanized and subjected to cardiac perfusion with PBS.

Liver samples were subsequently isolated and fixed with a 4% paraformaldehyde solution. Following three PBS washes, permeabilization was conducted using 2% PBST (2% Triton X-100, 0.05% sodium azide in PBS) for 3 days at RT. Samples were then rinsed with PBS and incubated in blocking buffer on an orbital shaker for 2 days at 4 °C. Immunostaining proceeded with F4/80 (Santa Cruz Biotechnology, clone A3- 1, 1:200) for 4 days at 4 °C. After incubation, samples were washed with washing buffer 3 times in RT and then overnight at 4 °C. Afterward, samples were then incubated with a secondary antibody (Life Technologies, 1:500) for 2 days at 4 °C and washed with washing buffer 3 times in RT then overnight at 4 °C. Following three PBS washes, nuclear staining was performed using DAPI for 1.5 h at RT. Subsequent PBS washing preceded tissue clearing with Rapiclear® reagent (SunJin Lab, cat: RC152001). Cleared specimens were mounted in Glass Bottom Cell Culture Dish (NEST) for imaging. Images were acquired using a confocal microscopy system(Leica, TSC SP8) with HC PL APO CS2 10×/0.40 DRY objective, with a scanning speed of 200 Hz and a step size of 2.0 μm. The X/Y acquiring format was set as 1024×1024, and Z-stack size was determined by specimen thickness respectively.

### Luciferase reporter assay

The *Pp6* 3’ UTR was cloned using primers: *Pp6* Forward, 5’-CCGCTCGAGCTC AAATGCTGCCTCTTGCCTTTTTTTTTAAT-3’, Reverse, 5’-ATAAGAATGCGG CCGCGAGGTTTACAGCCGGGTTGA-3’. The mutagenic primers used for *Pp6* were Mutant Forward, 5’-CCGCTCGAGCTCAAATGCTGCCGCGTACATTTTTT TTTAAT-3’. A genomic fragment corresponding to the *Pp6* 3’ UTR was cloned into the multiple cloning site of the psiCheck-2 synthetic firefly luciferase reporter plasmid (Promega, #C8021). Plasmid integrity was verified through DNA sequencing.

RAW264.7 cells were seeded in 48-well plates and transfected with a complex containing 100 ng 3’ UTR luciferase reporter vector and 50 pmol *miR-31* mimics or controls. Following 24-hour incubation post-transfection, cells lysis was performed. Luciferase activity quantification was conducted using the Dual-Luciferase Reporter Assay System (Promega) with measurements taken on a Lumat^3^ LB 9508 Single Tube Luminometer instrument (Berthold Technologies). Each experiment was performed in triplicate. The ratio of Renilla-to-Firefly luciferase was calculated for each well.

### Yeast-two hybrid screening

Sequence encoding mouse Pp6 was inserted into the pGBKT7 vector (BD-Pp6) and transformed into Y2H Gold yeast strain using Yeastmaker™ Yeast Transformation System 2 (Clontech, Cat. 630440). A normalized universal mouse cDNA library transformed into Y187 yeast strain was purchased from Clontech (Cat. 630482). Library screening was performed using Matchmaker® Gold Yeast Two-Hybrid System (Clontech, Cat. 630489) according to the manufacturer’ s instructions. Selected clones were isolated and sequenced. The prey plasmids were rescued and the positive interactions were confirmed using minimal media quadruple dropout plus X-α-Gal and Aureobasidin A (QDO/X/A).

### Western blotting

Cells lysis was performed using a buffer containing 50 mM Tris-HCl (pH 7.4), 5 mM sodium fluoride, 5 mM sodium pyrophosphate, 1 mM EDTA, 1 mM EGTA, 250 mM mannitol, 1% (v/v) Triton X-100, supplemented with protease inhibitor (Thermo Scientific) and phosphatase inhibitor A&B (Thermo Scientific). The proteins (10-30 μg) were loaded onto SDS-PAGE. Immunoblotting was performed with the following antibodies: Rabbit anti-PP6C (Merck, 1:1000), Rabbit anti-PFKFB1 (abcam, 1:500), Mouse anti-β-actin (Proteintech, 1:10000), HRP-labeled goat anti-mouse IgG (H + L) (Beyotime, 1:2000), and HRP-labeled goat anti-rabbit IgG (H + L) (Beyotime, 1:2000). Protein signals were visualized using Immobilon Western Chemiluminescent HRP Substrate (Millipore Sigma) and captured with GE Amersham Imager 600 (GE Healthcare).

### Unbiased metabolomics analysis

BMDMs isolated from six mice were cultivated. The samples were snap frozen and metabolites were extracted with 80% ice-cold methanol. Liquid chromatography-mass spectrometry (LC-MS) was performed on an Ultimate 3000-Velos Pro system equipped with a binary solvent delivery manager and a sample manager, coupled with a LTQ Orbitrap Mass Spectrometer equipped with an electrospray interface (ThermoFisher Scientific). The raw data were processed with progenesis QI (Walters Corporation) and Principal component analysis (PCA) and hierarchical clustering of metabolites were analyzed.

### Detection of NO production

Cells were lysed with cell lysis buffer for Western and IP (P1003, Beyotime) and the concentration of NO was determined using an NO assay kit (S0021, Beyotime) based on the Griess reaction. All samples were prepared according to the manufacturer’s protocol.

### Flow cytometry

BMDMs were initially digested using 0.5% trypsin-EDTA solution (Gibco) and were stained with Fc blocker and Live/dead Fixable Blue Dead Cell Stain Kit (Thermo Scientific cat. L34962) for 20 min at 4 ℃. Cells were then washed with PBS containing 2% FBS. Cell fixation and permeabilization were performed using the Cytosis/Cytoperm Fixation/Permeabilization Solution Kit (BD Biosciences cat. 554714). Cells stained with antibody AF-700 anti-hu/mo Arginase 1(Invitrogen, 1:200) for 30 min at 4 ℃. Samples were subsequent run on a BD Fortessa instrument and the data were analyzed using FlowJo software.

### RNA-seq and transcriptome analysis

BMDMs isolated from WT and *Pfkfb1*^-/-^ mice underwent two PBS washes prior to RNA extraction. Total RNA isolation was performed using RNAiso Plus (Takara Bio), followed by purification with magnetic oligo (dT) beads after denaturation. Purified mRNA samples were reverse-transcribed into fragmented DNA samples and adenylated at the 3′ ends. Library preparation involved adapter ligation, followed by DNA quantification using Qubit (Invitrogen). After cBot cluster generation, DNA samples were sequenced by an Illumina NovaSeq 6000 instrument from Shanghai Xu ran Biological Technology Co., LTD. Raw sequencing data were converted into FASTQ format, and transcript per million fragments mapped (fragments per kilobase) was calculated and log2-transformed using Cuffnorm. Differential gene transcripts were analyzed with DESeq, followed by gene set enrichment analysis (GSEA).

## Statistical analysis

All data were analyzed using GraphPad Prism version 8.0, and are presented as the mean ± SEM. Statistical comparisons employed Student’s t-test for parametric data and Log-rank test for survival analysis. The probability values of *P* < 0.05 were considered statistically significant, with the following notation scheme applied: \**P* < 0.05, \*\**P* < 0.01, \*\*\**P* < 0.001, and \*\*\*\**P* < 0.0001. Non-significant results were designated as “ns”. The error bars depict the SEM.

## Data availability

The RNA sequencing data generated from BMDMs are publicly accessible through the GEO database under accession code GSE291689. All source data supporting the findings described in this manuscript accompany the publication.

## Acknowledgments

The authors would like to thank Prof. X. Jia (Clinical Medicine Scientific and Technical Innovation Park, Shanghai Tenth Peoples Hospital) for P2 laboratory; Dr. L. Xia and Dr. L. Meng (Proteomics Core of College of Basic Medical Sciences, SJTU-SM) for protein LC-MS analyses.

## Author Contributions

**Conceptualization**: Honglin Wang, Li Fan, Yang Sun

**Data curation**: Li Fan, Yang Sun, Fangzhou Lou

**Formal analysis**: Li Fan, Yang Sun, Fangzhou Lou, Zilong Fang, Wengxiang Ding, Xiangxiao Li, Yan Li, Qingqing Shen, Siyu Deng, Jihuan Liang, Fengjiao Zhang, Sibei Tang, Zhikai Wang, Xiaojie Cai, Jiajia Tong, Zhenyao Xu, Jing Zou, Qing Yang, Honglin Wang

**Investigation**: Li Fan, Yang Sun **Supervision**: Honglin Wang **Writing – original draft**: Li Fan

**Writing – review & editing**: Yang Sun, Honglin Wang

---

**S1 Fig.**
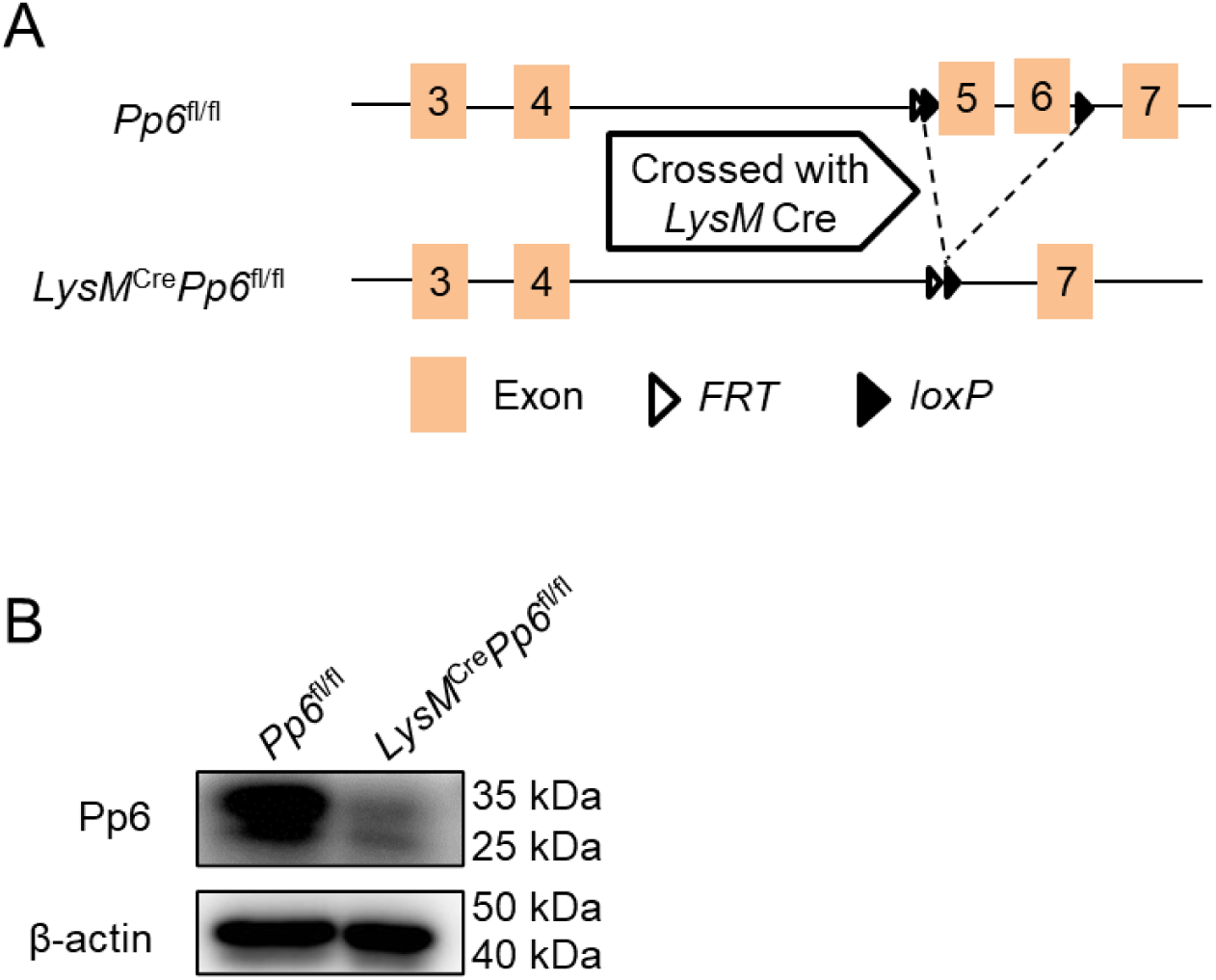
Strategies for *Pp6* conditional knockout mouse construction **(A-B)** Schematic of *Pp6* deletion in *LysM*^Cre^*Pp6*^fl/fl^ mice (A) and western blot analysis of Pp6 expression in BMDMs from *LysM*^Cre^*Pp6*^fl/fl^ and *Pp6*^fl/fl^ mice (B).

**S2 Fig.**
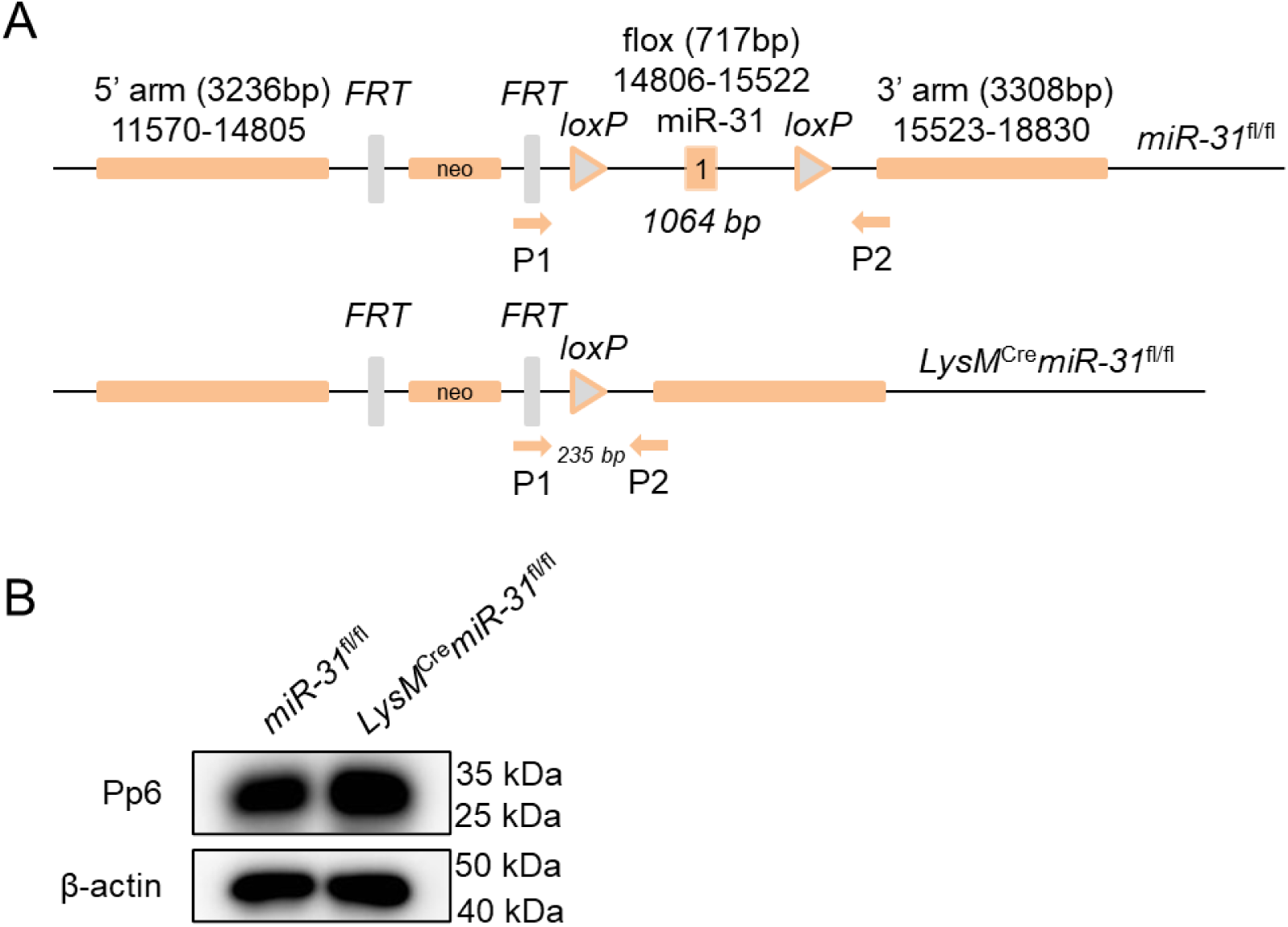
Strategies for *miR-31* conditional knockout mouse construction **(A-B)** Schematic of *miR-31* deletion in *LysM*^Cre^*miR-31*^fl/fl^ mice (A) and western blot analysis of Pp6 expression in BMDMs from *LysM*^Cre^*miR-31*^fl/fl^ and *miR-31*^fl/fl^ mice (B).

**S3 Fig.**
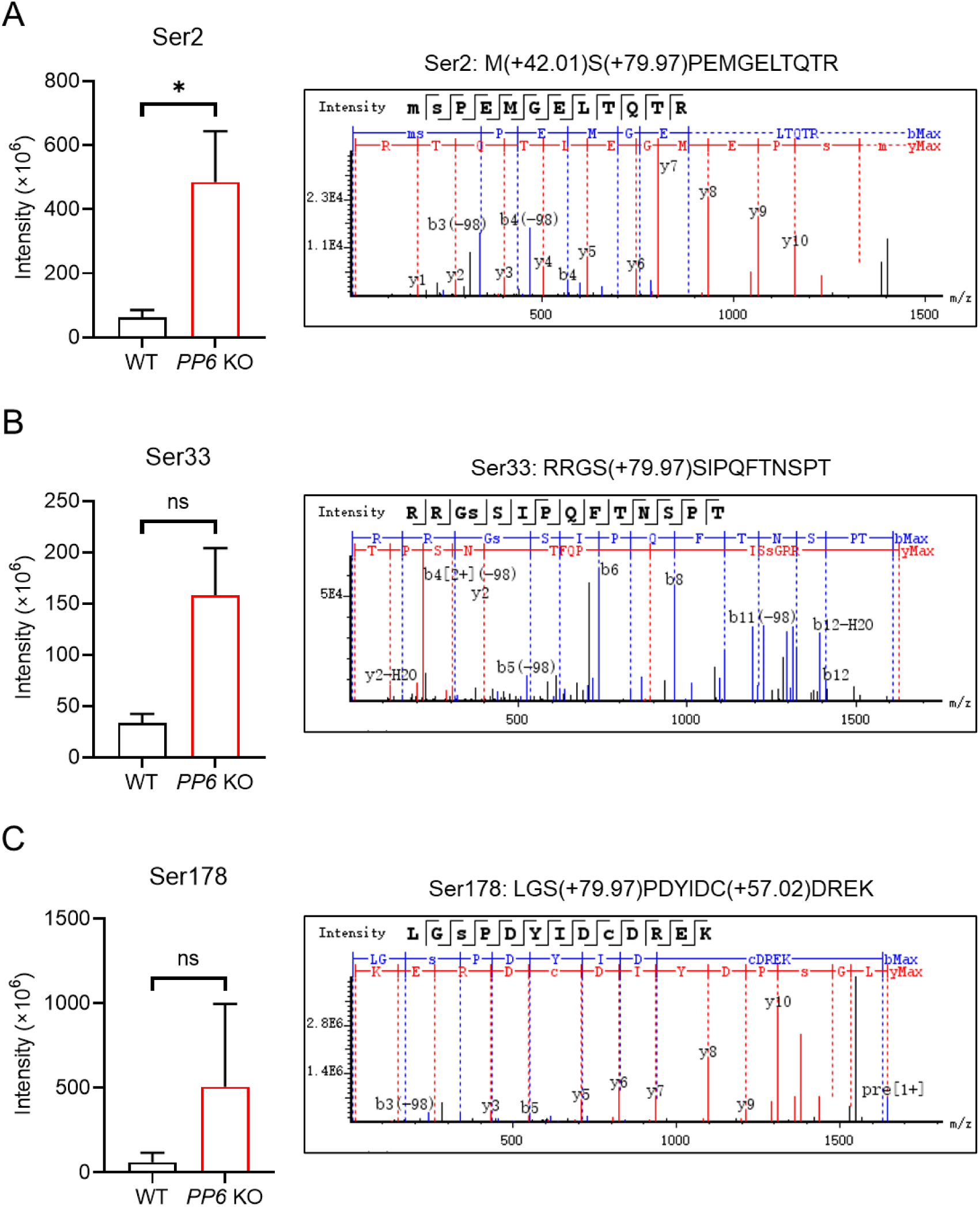
Ser-2, Ser-33 and Ser-178 sites were hyper-phosphorylated in *PP6*- deficient 293FT cells **(A-C)** LC-MS/MS-based detection of phosphorylation sites on PFKFB1 immunoprecipitates from WT and *PP6*-deficient 293FT cells. Left: Intensity of Ser-2 (A), Ser-33 (B), Ser-178 (C) in WT and *PP6*-deficient 293FT cells. *P*-values were determined by two-tailed unpaired *t*-test (mean ± SEM). ns: not significant, \**P* < 0.05. Right: Representative mass spectrum of Ser-2 (A), Ser-33 (B), Ser-178 (C).

**S4 Fig.**
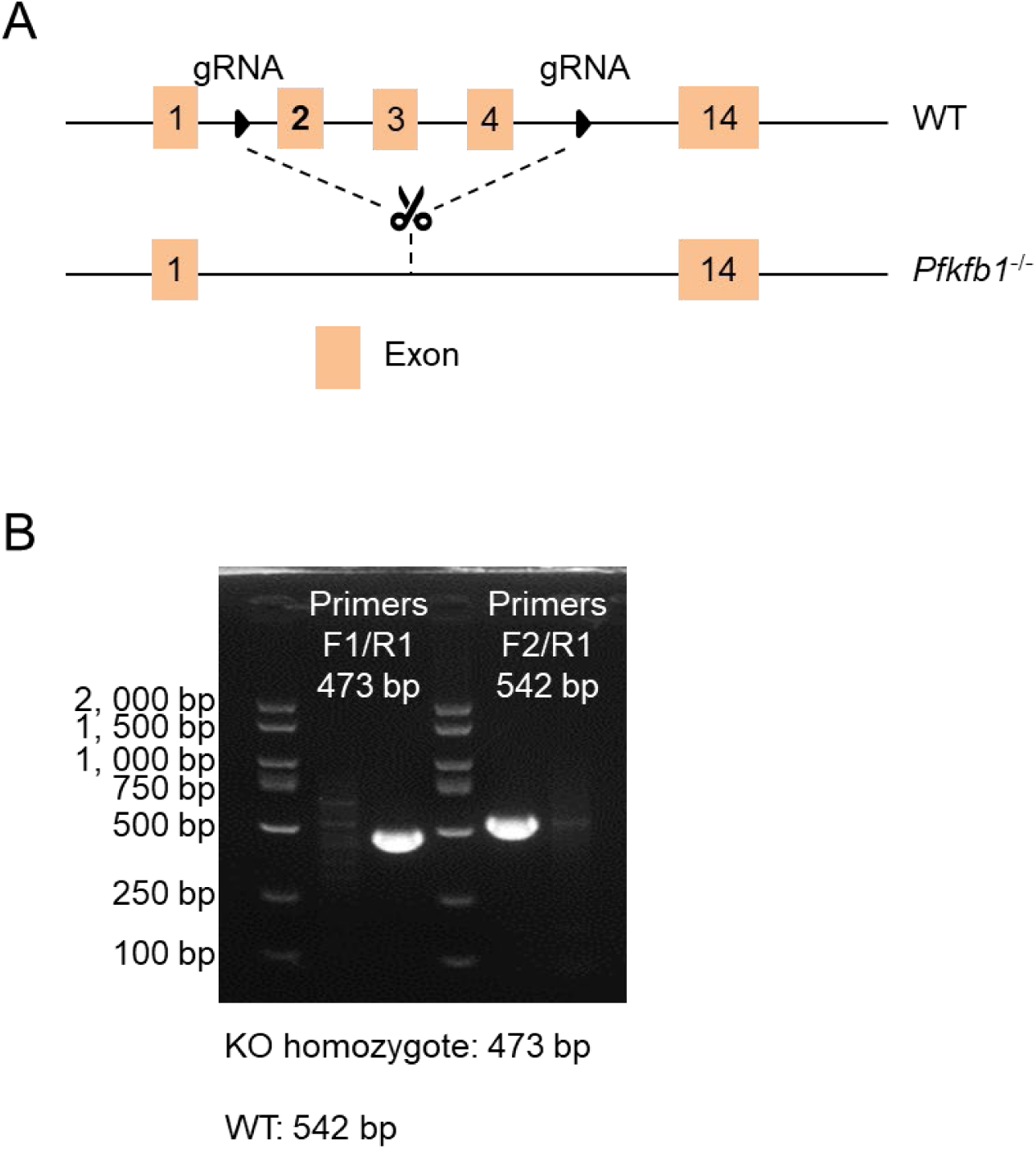
Strategies for *Pfkfb1* knockout mouse construction **(A-B)** Schematic of *Pfkfb1* deletion in *Pfkfb1*^-/-^ mice (A) and *Pfkfb1* genotyping of the toes derived from WT and *Pfkfb1*^-/-^ mice (B).

**S1 Table.**
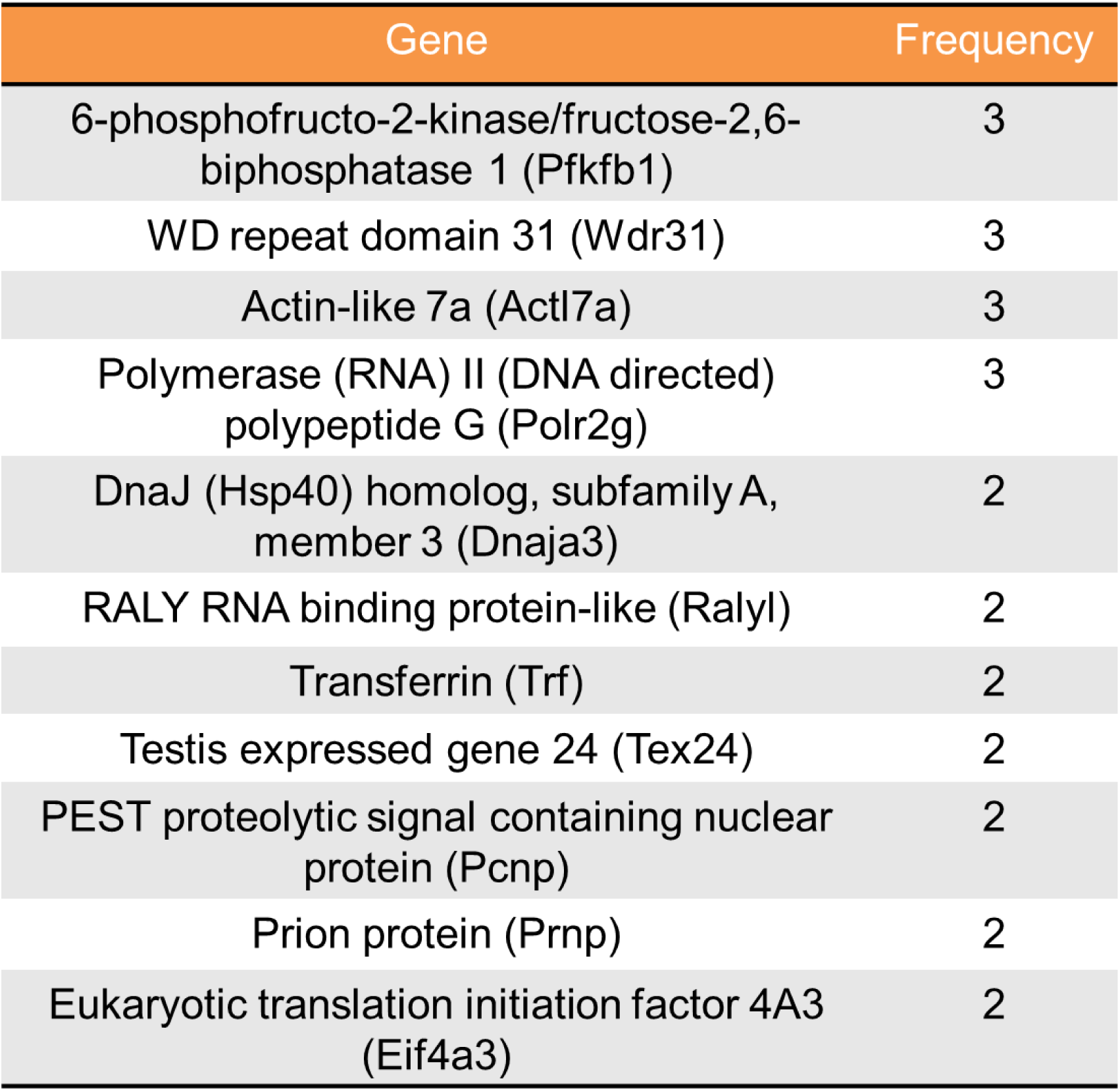
Top 11 genes in yeast two-hybrid screening.

